# A combination of two human neutralizing antibodies prevents SARS-CoV-2 infection in rhesus macaques

**DOI:** 10.1101/2021.09.27.462074

**Authors:** Ronald R. Cobb, Joseph Nkolola, Pavlo Gilchuk, Abishek Chandrashekar, Robert V. House, Christopher G. Earnhart, Nicole M. Dorsey, Svetlana A. Hopkins, Doris M. Snow, Rita E. Chen, Laura A. VanBlargan, Manuel Hechenblaikner, Brian Hoppe, Laura Collins, Milan T. Tomic, Genevieve H. Nonet, Kyal Hackett, James C. Slaughter, Michael S. Diamond, Robert H. Carnahan, Dan H. Barouch, James E. Crowe

**Author notes:** Correspondence: Dan Barouch,; James Crowe,. These authors contributed equally (Ronald R. Cobb, Joseph Nkolola, Pavlo Gilchuk).

## Abstract

Human monoclonal antibody (mAb) treatments are promising for COVID-19 prevention, post-exposure prophylaxis, or therapy. However, the titer of neutralizing antibodies required for protection against SARS-CoV-2 infection remains poorly characterized. We previously described two potently neutralizing mAbs COV2-2130 and COV2-2381 targeting non-overlapping epitopes on the receptor-binding domain of SARS-CoV-2 spike protein. Here, we engineered the Fc-region of these mAbs with mutations to extend their persistence in humans and reduce interactions with Fc gamma receptors. Passive transfer of individual or combinations of the two antibodies (designated ADM03820) given prophylactically by intravenous or intramuscular route conferred virological protection in a non-human primate (NHP) model of SARS-CoV-2 infection, and ADM03820 potently neutralized SARS-CoV-2 variants of concern *in vitro*. We defined 6,000 as a protective serum neutralizing antibody titer in NHPs against infection for passively transferred human mAbs that acted by direct viral neutralization, which corresponded to a concentration of 20 μg/mL of circulating mAb.

## INTRODUCTION

In the past decades, two pathogenic human coronaviruses, severe acute respiratory syndrome (SARS-CoV) and Middle East respiratory syndrome CoV (MERS-CoV), have been reported to cause severe respiratory tract disease associated with high morbidity and mortality. In December 2019, the severe acute respiratory syndrome coronavirus 2 (SARS-CoV-2) emerged in Wuhan, Hubei province, China (Wang et al., 2020). SARS-CoV-2 is the causative agent of the current worldwide COVID-19 outbreak. The pandemic caused by COVID-19 has made the development of countermeasures an urgent global priority (Chan et al., 2020; Chen et al., 2020a; Li et al., 2020; Wu et al., 2020a; Zhou et al., 2020). Safe and effective vaccines and therapeutics are essential to combat this global pandemic.

Initial work identified that SARS-CoV-2 uses the angiotensin-converting enzyme 2 (ACE2) protein from bats, civet cats, swine, non-human primates, or humans as an attachment and entry receptor (Letko et al., 2020; Wan et al., 2020; Zhou et al., 2020). As with related coronaviruses, interaction with ACE2 is mediated principally through the viral spike (S) protein. Hence, S on the surface of the virion is the main target for neutralizing antibodies on these coronaviruses. This homotrimeric glycoprotein is anchored in the viral membrane and consists of two subunits, S1, containing the N-terminal domain (NTD) and host cell receptor binding domain (RBD), and S2, which contains the fusion peptide (Walls et al., 2020; Wrapp et al., 2020). The S protein RBD directly interacts with the peptidase domain of ACE2 (Letko et al., 2020; Wan et al., 2020; Wrapp et al., 2020; Zhou et al., 2020). Recent studies of the S protein structure have shown that the protein exists in different conformations (Cai et al., 2020; Walls et al., 2020). Initially, the RBD switches from a closed conformation to an open conformation to allow hACE2 interaction. Upon interaction with the hACE2 receptor and TMPRSS2 priming, S2 undergoes a dramatic conformational change to trigger host membrane fusion (Fan et al., 2020).

The RBD is the primary target of most potently neutralizing anti-SARS-CoV-2 antibodies identified to date (Cao et al., 2020; Ju et al., 2020; Pinto et al., 2020; Rogers et al., 2020; Shi et al., 2020; Wu et al., 2020b; Zost et al., 2020a). The RBD is also the main antigenic site for neutralizing antibody responses in current and experimental COVID-19 vaccines (Chen et al., 2020b; Mulligan et al., 2020; Zang et al., 2020). Previous studies established a non-human primate (NHP) model for SARS-COV-2 infection (Chandrashekar et al., 2020; Yu et al., 2020) demonstrating protection from viral infection by transfer of a high-dose of ACE2-blocking monoclonal antibodies (Zost et al., 2020a). Currently available antibody therapeutics that have received EUA from the FDA were approved for post-exposure treatment, not for pre-exposure prophylaxis (FDA, 2020, 2021a, b). Prophylaxis with passive antibody therapy could be important as an option for individuals at high risk of disease from SARS-CoV-2 infection who cannot be adequately vaccinated, including immunocompromised individuals or others who respond poorly to vaccination (AstraZeneca, 2021; Loo et al., 2021).

Here, we evaluated the prophylactic efficacy of low or moderate doses of two different human mAbs targeting non-overlapping neutralization epitopes in the RBD domain (Zost et al., 2020a; Zost et al., 2020b), which we assessed individually or in combination. It has been previously shown that antibody cocktails can limit the risk of viral mutations that escape antibody neutralization more efficiently than monotherapy (Baum et al., 2020b; Chen *et al*., 2021b; Greaney et al., 2021). The antibody COV2-2381 binds directly to the receptor-binding motif on the RBD on an S protomer in the open position. In contrast, the antibody COV2-2130 binds a non-overlapping site on the RBD that is accessible in either the open or closed S protomer conformation. We engineered the Fc portion of these antibodies to contain mutations that extend half-life (M252**<u>Y</u>**/S254**T**/T256**E**, designated YTE) (Richards et al., 1999; Uppal et al., 2015; Wang et al., 2015) and also to reduce Fcγ receptor binding (L234A/L235A, designated LALA) (Lund et al., 1991; Wines et al., 2000; Xu et al., 2000). One conceptual advantage of this approach is that the use of these antibodies lacking Fc-mediated effects allowed us to assess the level of serum neutralizing activity needed *in vivo* to achieve efficacy in the absence of confounding variables. The resulting recombinant mAbs were designated mAb COV2-2130-YTE-LALA and mAb COV2-2381-YTE-LALA, and a two-mAb cocktail that is a 1:1 mixture of the two was designated ADM03820. The results demonstrate that ADM03820 protects against challenge with SARS-CoV-2 in the lungs and nasopharynx in a dose-dependent manner, and define titers of passively-transferred neutralizing antibodies that are necessary for protection in NHPs. In addition, our results support the use of antibody cocktail that could be administered by either intravenous or intramuscular route and that neutralizes SARS-CoV-2 variants of concern. This work provides evidence for developing a cocktail of antibodies as prophylaxis against SARS-CoV-2 in high-risk individuals.

## RESULTS

### ADM03820 antibody cocktail is detected at primary sites of SARS-CoV-2 infection in NHPs when administered by IV or IM routes (Study 1)

In this study, we used a rhesus macaque SARS-CoV-2 challenge model for pre-clinical development studies of a prophylactic cocktail ADM03820 comprising two engineered mAbs, COV2-2130-YTE-LALA and COV2-2381-YTE-LALA. We first assessed the human antibody concentration in serum and at primary sites of infection (*e.g*., upper and lower respiratory tract mucosa) after 11.7 mg/kg intramuscular (IM) or 31.3 mg/kg intravenous (IV) administration of ADM03820 in rhesus monkeys (**Figure 1A**). Circulating human mAbs were detected at high levels in serum on day 0 after administration (median 193 μg/mL after IM or 520 μg/mL after IV administration) and persisted in serum for >80 days, exhibiting a slow and gradual decline. The median human IgG serum concentration was 9 μg/mL on day 84 after IM or 26 μg/mL after IV administration (**Figure 1B**). Notably, ADM03820 antibodies also were detected in respiratory tract secretions, including bronchoalveolar lavage (BAL) and nasopharyngeal (NP) swabs up to 60 days after administration and at concentrations ranging from 10 ng/mL (the assay limit of detection) to 270 ng/mL (**Figure 1C,D**). The concentration of human antibodies in these secretions *in vivo* prior to collection is expected to be higher, given that specimen collection from the mucosa sites with saline washes resulted in antibody dilution.

**Figure 1.**
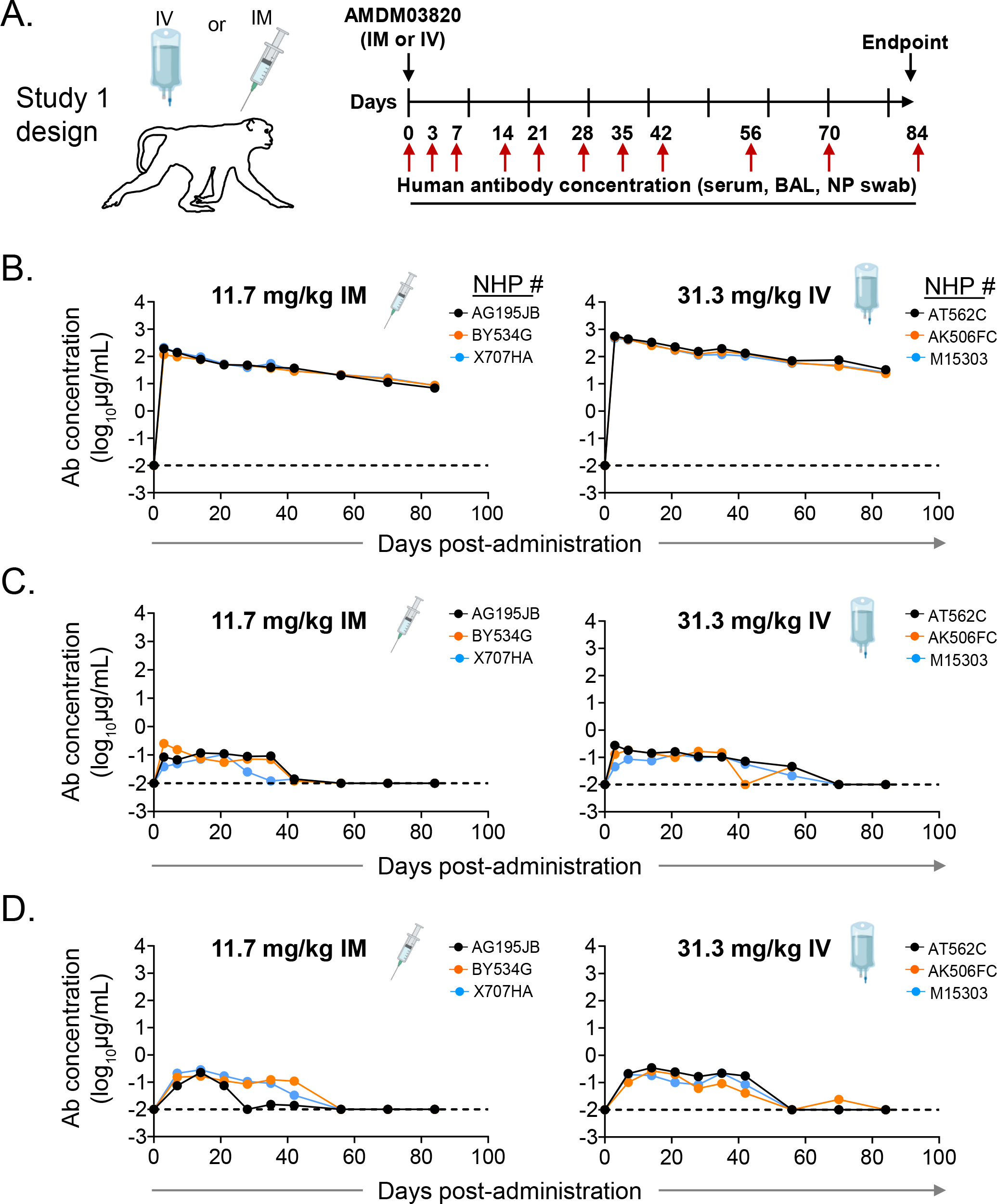
Pharmacokinetics and biodistribution of ADM03820. (**A**) Schema of study design. Different doses of antibody cocktail ADM03820 (containing COV2-2130/YTE-LALA and COV2-2381/YTE-LALA at a 1:1 ratio) were administered to rhesus monkeys (n=3 per group) by IV (11.7 or 31.3 mg/kg) or IM (11.7 or 31.3 mg/kg) route. Human antibody concentration was assessed by ELISA in (**B**) serum, (**C**) BAL, or (**D**) nasal swab eluate samples at indicated time points after ADM03820 administration. The dotted horizontal line depicts the assay limit of detection.

### ADM03820 antibody cocktail potently neutralizes variants of concern

ADM03820 exhibited broad and potent neutralizing activity *in vitro* with half-maximal inhibitory concentration values <25 ng/mL, including potent neutralization of viruses representing wild-type SARS-CoV-2 WA1/2020 with or without D614G mutation, authentic B.1.1.7 virus, authentic B.1.617.2 virus, and chimeric Wash-B.1.351 and Wash-B.1.1.28 viruses, which contain an S gene from B.1.351 or B.1.1.28, respectively, in the backbone of WA1/2020 (Chen *et al*., 2021a) (**Table 1**). Collectively, these results showed prolonged persistence of administered human antibodies in serum and respiratory mucosa at concentrations sufficient for neutralization of currently circulating viral variants.

**Table 1.** Neutralization breadth of ADM03820 against SARS-CoV-2 variants of concern. [IC_50_ (ng/mL) against indicated virus]^2^ ^1^Neutralizing activity of ADM03820 against authentic SARS-CoV-2 WA1/2020, authentic SARS-CoV-2 WA1/2020 bearing D614G mutation, or authentic B1.1.7, authentic B.1.617.2, chimeric Wash-B1.351, and chimeric Wash-B.1.1.28 viruses was assessed using a focus reduction neutralization test (FRNT). ^2^Half-maximal inhibitory concentration (IC50) values are shown and represent the average of technical duplicates and two independent experiments.

| **Table 1. Neutralization breadth of ADM03820 against SARS-CoV-2 variants of concern<sup>1</sup>**<br>[IC<sub>50</sub> (ng/mL) against indicated virus]<sup>2</sup> |  |  |  |  |  |
| --- | --- | --- | --- | --- | --- |
| **WA1/2020** | **D614G** | **B.1.1.7 (Alpha)** | **Wash-B1.1.351 (Beta)** | **B.1.617.2 (Delta)** | **Wash-B.1.1.28 (Gamma)** |
| 28 | 21 | 20 | 19 | 25 | 8 |

### Protective efficacy of ADM03820 in nonhuman primates (Study 2)

To evaluate the protective efficacy of ADM03820, animals received various doses of the ADM03820 by either IM or IV route followed by a viral challenge with 10^5^ tissue culture infectious dose (TCID_50_) 3 days later (**Figure 2A**). We then measured the circulating human antibody concentration in serum and serum neutralizing titers up to day 14 following IM or IV administration. While antibody concentration was below the limit of detection in the sham-treated group, animals in the antibody-treatment groups exhibited mAb levels proportional to the dose and route of administration of the combination product (**Figure 2B**). The antibody concentration in serum peaked approximately three days post-administration and remained constant throughout the remaining 14 days of the study.

**Figure 2.**
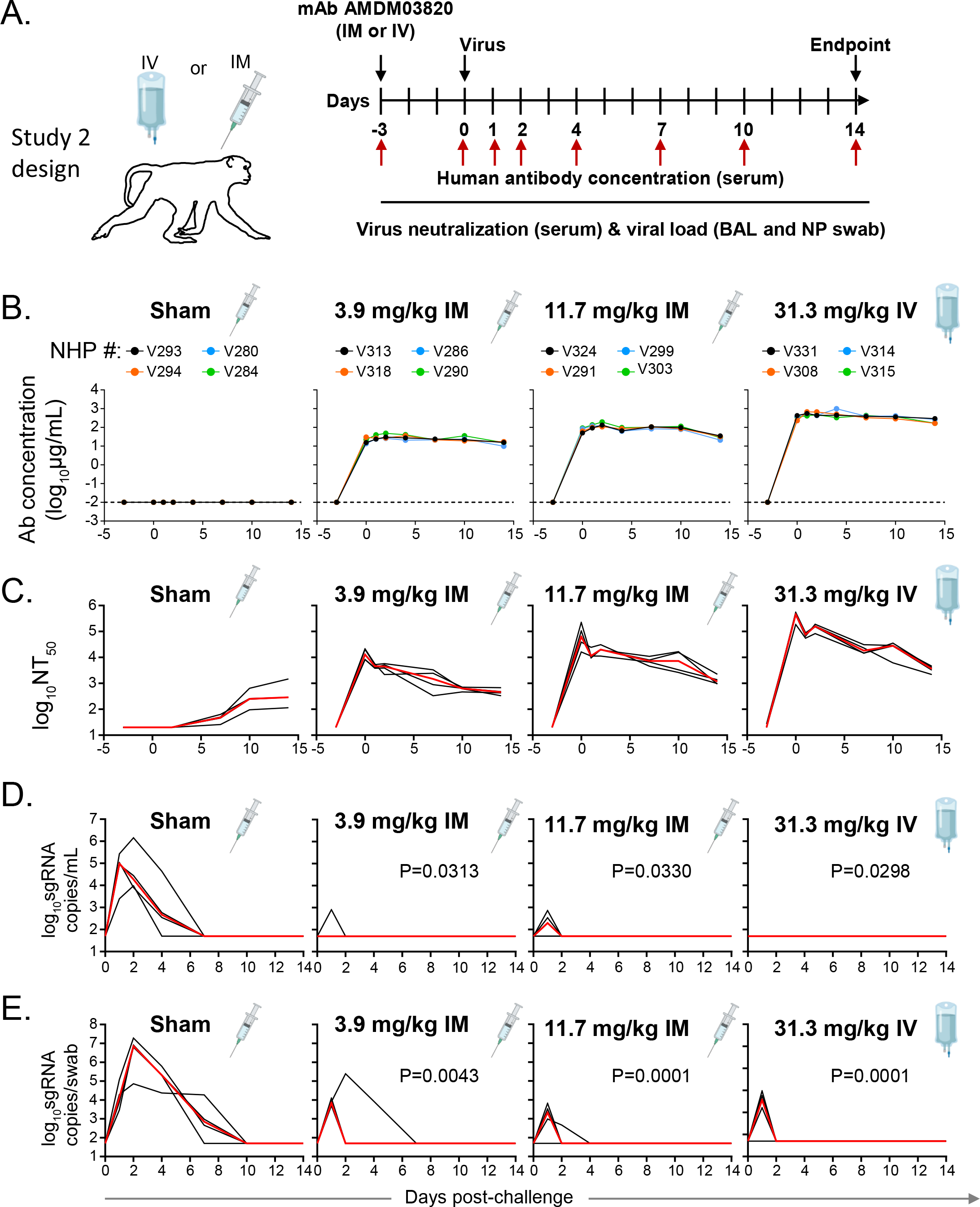
Pharmacokinetics, antibody neutralizing titers, and prophylactic efficacy of ADM03820 mAbs in SARS-CoV-2-challenged NHPs. (**A**) Schema of study design. Different doses of ADM03820 were administered to rhesus monkeys (day -3) by IM (3.9 or 11.7 mg/kg) or IV (31.3 mg/kg) route (n=4 per group). One group of NHPs was left untreated (sham; n=4) and served as a control. Animals in all groups were challenged with 10^5^ TCID_50_ SARS-CoV-2 by the intranasal and intratracheal routes on day 0. (**B**) Human antibody concentration in serum was assessed by ELISA at indicated time points after ADM03820 administration and viral challenge. (**C**) Total neutralizing antibody titers were assessed in serum at indicated time points using pseudovirus neutralization assay. The red line indicates the median titer of neutralizing antibodies in each group. (**D**) Subgenomic RNA (sgRNA) levels were assessed at various time points after viral challenge in bronchoalveolar lavage (BAL) samples using RT-qPCR. (**E**) Subgenomic RNA (sgRNA) levels were assessed at various time points after viral challenge in nasopharyngeal (NP) swab samples. Each black curve shows the measurements from individual animals, with red lines indicating the median values of measurements for animals within each treatment group. Neutralization assay limit of detection = 50 copies/mL or 50 copies/swab for panels (D) and (E). For statistical analysis, refer to *Methods* section.

We observed high circulating neutralizing antibody titers by pseudovirus neutralization assays in all ADM03820 treatment groups but not in the sham-treated control group. However, sham-treated control animals developed low-level neutralizing titers beginning around day 6, presumably due to the induction of natural host immunity (**Figure 2C**). In general, the overall neutralizing antibody titers were consistent with the pharmacokinetic data for the same treatment groups.

We assessed the kinetics of viral loads up to day 14 following viral challenge in BAL and NP swab samples by determining the levels of SARS-CoV-2 sub-genomic RNA (sgRNA), which distinguishes replicating virus from input challenge virus, using reverse-transcriptase-polymerase chain reaction (RT-PCR) (Chandrashekar et al., 2020; Wolfel et al., 2020; Yu et al., 2020). High levels of sgRNA were observed in the sham controls (**Figure 2D-E**) with a median peak of 5.0 (range = 3.3 to 5.4) log_10_ sgRNA copies/mL in BAL fluid and 6.9 (range = 4.9 to 7.3) log_10_ copies per swab of sgRNA in NP. As expected, peak viral loads occurred between days 1 to 4 after challenge. All treatment groups showed nearly full protection from viral replication in the BAL fluid, although individual animals displayed low-level, transient viral replication on day 1, which was eliminated by day 2 (**Figure 2D**). Although somewhat higher sgRNA levels were observed in some animals in the NP swabs on day 1, similar to BAL fluid, most treated animals quickly eliminated detectable virus by day 2 (**Figure 2E**), with the exception of one animal in the group receiving the lowest dose (3.9 mg/kg IM) and one animal in the group receiving 11.7 mg/kg dose.

### Protective efficacy of individual mAbs of the cocktail in nonhuman primates (Study 3)

The next challenge study was conducted after prophylactic administration of the individual 2130-YTE-LALA or 2381-YTE-LALA antibodies and was followed by quantitative serum antibody levels and virologic protection measurements as in the challenge study above (**Figure 3A**). As expected, the concentration of circulating human antibodies was below the level of detection in the sham-treated group. In contrast, animals that received either mAb demonstrated concentrations in serum proportional to the administered dose (**Figure 3B**). Peak antibody concentration was observed within three days of administration and remained constant throughout the study. Serum neutralizing titers of administered individual mAbs showed similar peak and kinetics to those seen with the ADM03820 cocktail (**Figure 3C**). Sham-treated animals showed low levels of neutralizing antibody activity by day six due to the host immune response to SARS-CoV-2 infection (**Figure 3C**).

**Figure 3.**
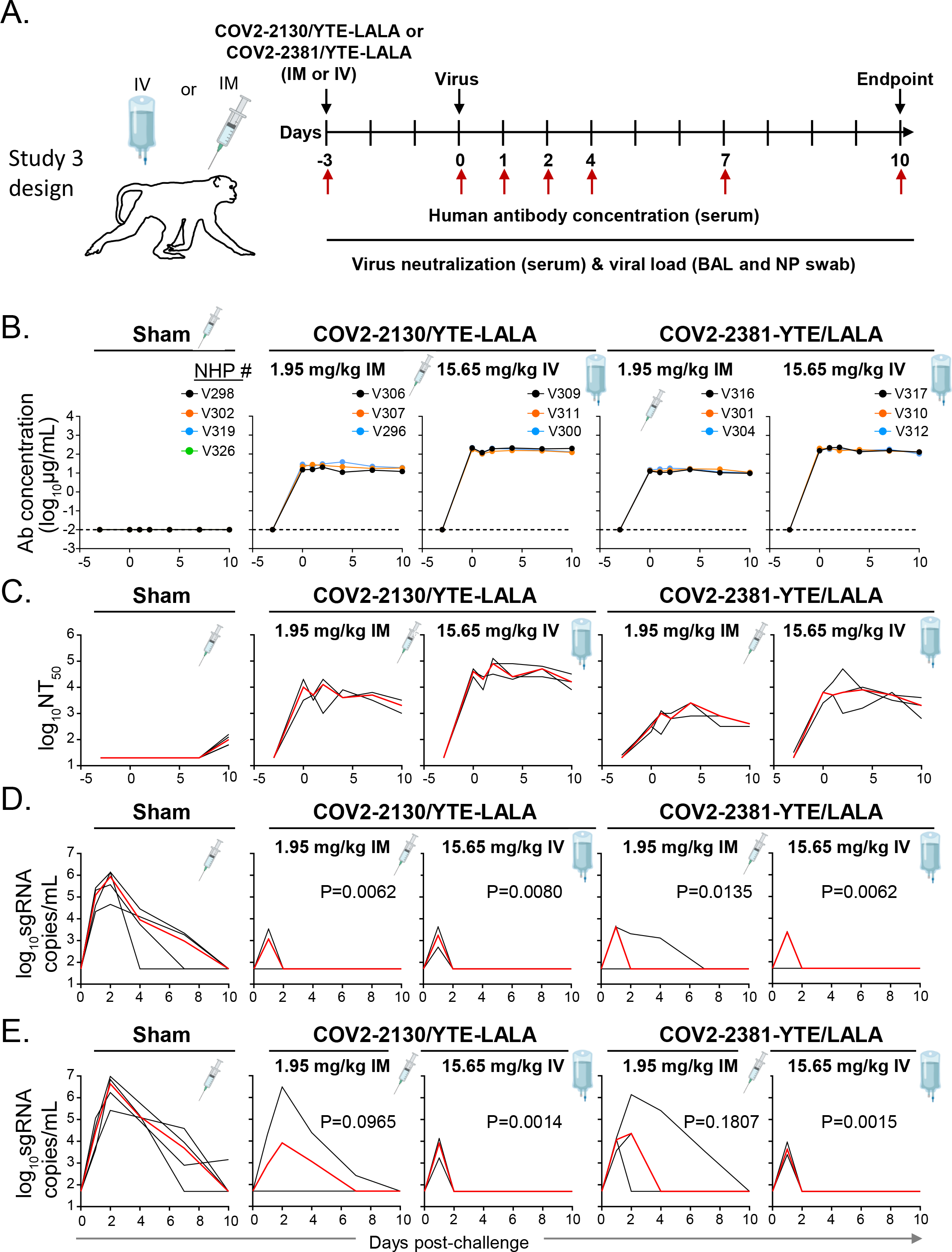
Pharmacokinetics, neutralizing titers, and prophylactic efficacy of individual mAbs of the cocktail in SARS-CoV-2-challenged NHPs. (**A**) Schema of study design. Individual mAbs COV2-2130/YTE-LALA or COV2-2381/YTE-LALA (n=3 NHP per group) were administered to rhesus monkeys (day -3) at different doses (1.95 mg/kg or 15.65 mg/kg) and routes (IM or IV) as indicated. One group of NHPs was left untreated (sham; n=4) to serve as controls. Animals in all groups were challenged with SARS-CoV-2 by the intranasal and intratracheal routes on day 0. (**B**) Human antibody concentration was assessed by ELISA in serum at indicated time points after indicated mAb administration and viral challenge. (**C**) Total neutralizing antibody titers were assessed in serum at indicated time points using a pseudovirus neutralization assay. Each black curve shows the measurements from an individual animal, with red lines indicating the median values of measurements for animals within each treatment group. (**C**) sgRNA levels were assessed after viral challenge at various time points in BAL samples using RT-qPCR. (**D**) sgRNA levels were assessed after viral challenge at various time points in nasopharyngeal swab samples. The red line depicts the median levels of sgRNA in each group. Each black curve shows an individual animal’s measurements, with red lines indicating the median values of measurements for animals within each treatment group. Neutralization assay limit of detection = 50 copies/mL or 50 copies/swab. For statistical analysis, refer to *Methods* section.

As evidenced by sgRNA levels, viral infection again was observed in all the sham-treated control animals in both BAL fluid and NP samples (**Figure 3D,E**). For both treatment mAbs, most animals quickly cleared virus by day two post-challenge after transient viral replication regardless of dose or route of administration, except for one animal in the 1.95 mg/mL 2381-YTE/LALA IM group (**Figure 3D**).

Similar levels of viral protection were observed in NP samples with the 15.65 mg/mL dose of either individual antibody (**Figure 3E**) as was observed with similarly high tested doses of the ADM03820 cocktail (**Figure 2E**). However, higher median viral loads were observed in the NP samples for both antibody treatments at the low dose of 1.95 mg/mL. This dose is two-fold lower than the lowest dose tested for the cocktail and likely represents viral breakthrough due to insufficient neutralizing antibody levels.

### Protective efficacy of ADM03820 that administered by IM route at low doses (Study 4)

To determine the minimally protective dose of the ADM03820 cocktail, animals were treated with two-fold decreasing doses of the antibody cocktail across four treatment groups from 3.91 mg/kg to 0.49 mg/kg by the IM route (**Figure 4A**). Circulating human antibody titers were not present in sham-treated animals and were consistent with the administered dose in the treatment groups (**Figure 4B**). The serum neutralizing antibody titer decrease was proportional to the administered ADM03820 dose and was observed across all four treatment groups but not observed in sham group animals (**Figure 4C**). BAL fluid viral load measurement suggested protection in the lower airways at all tested antibody doses, including at the 0.49 mg/kg dose (**Figure 4D**). However, increases in NP swab viral loads were seen across decreasing dose conditions, with no protection observed in the 0.98 mg/mL or 0.49 mg/mL groups (**Figure 4E**). These results suggested that a higher antibody dose would be necessary to control viral replication in the upper airways following IM administration.

**Figure 4.**
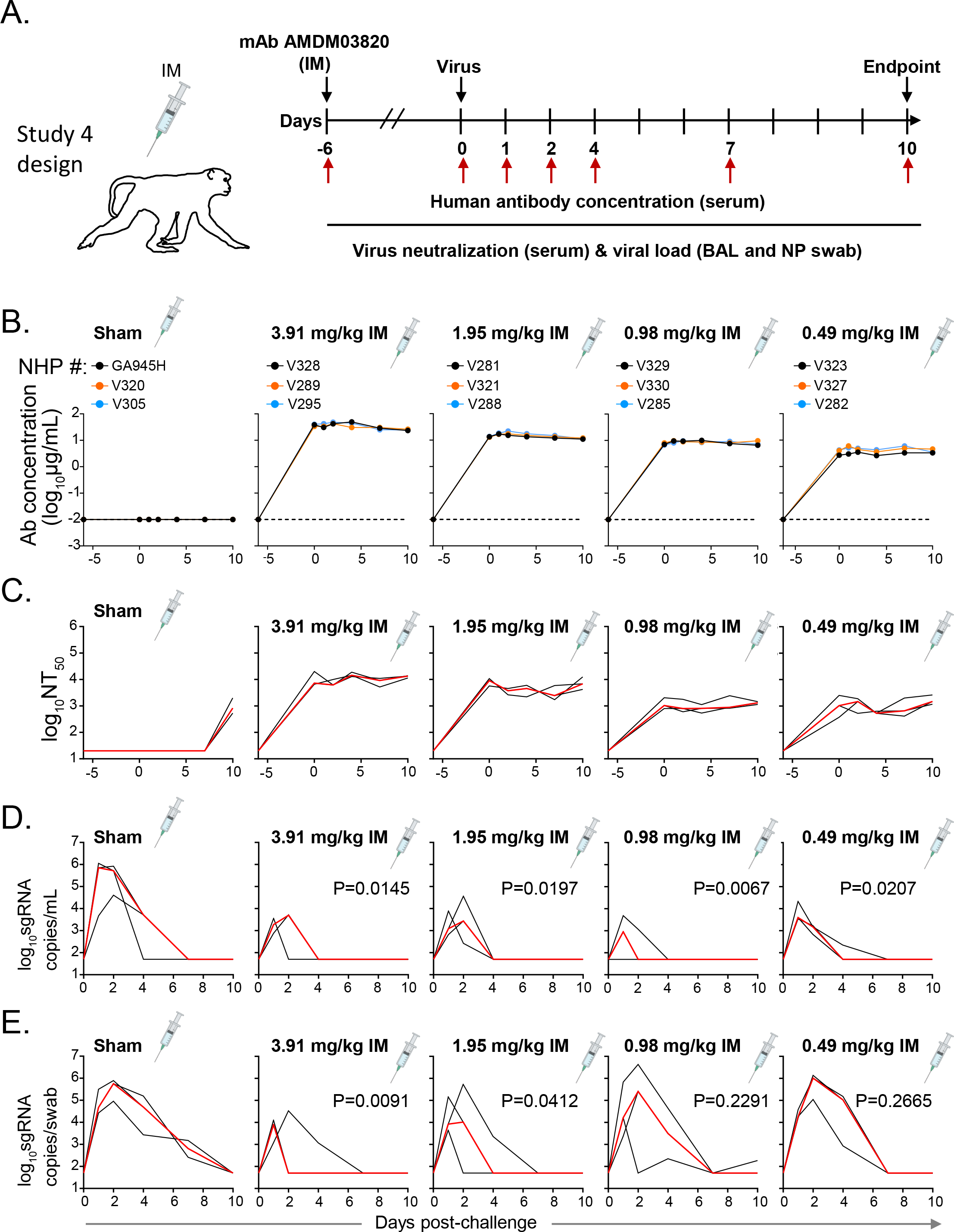
Pharmacokinetics, neutralizing titers, and prophylactic efficacy of ADM03820 in a dose de-escalation study and IM antibody administration in NHPs. (**A**) Schema of study design. Different doses of ADM03820 were administered to rhesus monkeys (day -6) by IM route (3.91, 1.95, 0.98, and 0.49 mg/kg; n=3 NHP per group). One group of NHPs was left untreated (sham; n=3) and served as a control. Animals in all groups were challenged with SARS-CoV-2 by the intranasal and intratracheal routes at day 0. (**B**) Human antibody concentration was assessed by ELISA in serum at indicated time points after ADM03820 administration and viral challenge. (**C**) Total neutralizing antibody titers in serum were assessed at indicated time points using a pseudovirus neutralization assay. The red line shows median titer of neutralizing antibodies in each group. (**D**) sgRNA levels were assessed at various time points after viral challenge in BAL samples using RT-qPCR. (**E**) sgRNA levels were assessed at various time points after viral challenge in nasopharyngeal swab samples. The red line depicts the median levels of sgRNA in each group. Each black curve shows measurements from an individual animal, with red lines indicating the median values of measurements for animals within each treatment group. Assay limit of detection = 50 copies/mL or 50 copies/swab. For statistical analysis, refer to *Methods* section.

### Defining protective serum antibody concentration and neutralizing antibody titer in NHP SARS-CoV-2 challenge model

We next estimated a protective threshold for prophylaxis with potent YTE-LALA Fc-region engineered human Abs that acted principally via direct virus neutralization *in vivo*. We performed an overall analysis using data from challenge studies 2, 3 and 4 above by comparing human mAb concentration in serum or half-maximal neutralizing titer values at the time of challenge to the time-weighted average values for the change of sgRNA viral load in BAL fluid or NP swabs from day 1 to day 10 after viral challenge (see *Methods* and **Table S1-2**). A threshold for virological protection in BAL fluid and NP was estimated to be equal or higher than 20 μg/mL for circulating human antibody concentration and equal to or higher than 6,000 for serum neutralizing antibody titer (NT_50_) (**Figure 5A-D**). Antibody levels above these thresholds conferred full protection in 83% to 93% of challenged NHP, which contrasted with 17% to 50% fully protected animals with antibody levels below these estimated protective thresholds (**Figure 5E**). Overall, our results suggested that high prophylaxis efficacy can be achieved with the cocktail of two YTE-LALA Fc-region engineered human Abs formulated as a cocktail ADM03820 and demonstrated the potential for IM delivery of human antibody-based therapeutics for COVID-19.

**Figure 5.**
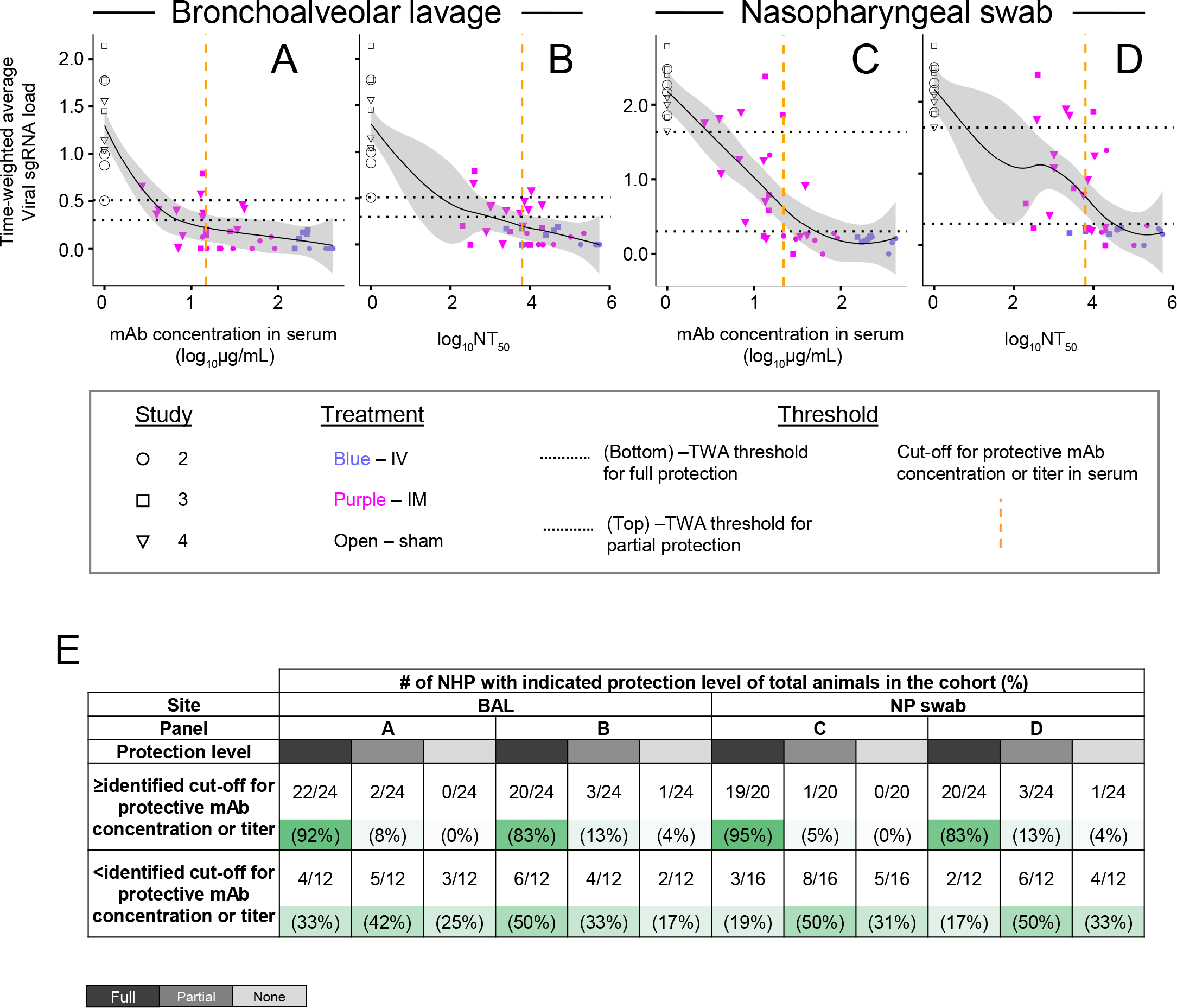
Human antibody concentration and antibody neutralizing titer in NHP serum associated with protection against viral challenge in BAL or NP samples. (**A-D**) The time-weighted average (TWA) values for the change of sgRNA viral load in BAL or NP swabs from day 1 to day 10 after viral challenge were compared to antibody concentration in serum or serum NT_50_ value for each animal from studies 2, 3 and 4 described in Figures 2 through 7. The fitting curves were estimated using Lowess curve smoothing method and are shown in black, and grey shading indicates the confidence interval. Shapes indicate individual animals, colors indicate route of antibody treatment, and animals from separate studies are shown with different shapes as detailed in the figure. Horizontal black dotted lines indicate designated TWA thresholds for full (bottom line) and partial (top line) protection. Vertical dotted orange dashed line in the graphs indicates designated estimated optimal cut-off for protective antibody concentration or titer in NHP serum. For calculation of TWA and cut-off values, refer to *Methods* section. (**E**) Percent animals that fully protected, partially protected, or non-protected determined using the estimated thresholds for protection as in panels **A-D**. Gradient of green shading visualize % of protected animals in which dark green indicates higher % of protected animals and light green indicates lower % of protected animals for each described condition.

## DISCUSSION

These studies provide insights into both quantitative and qualitative aspects of the use of human mAbs as medical countermeasures for COVID-19. First, we demonstrate the principle that prophylaxis against infection in NHPs can be achieved using neutralizing antibodies engineered to lack Fc-mediated functions. These data extend previous findings that demonstrated prophylaxis efficacy for neutralizing mAbs with intact Fc-mediated functions in NHPs (Baum et al., 2020a; Jones et al., 2021; Zost et al., 2020a; Winkler et al., 2021). Second, the data show excellent protection by antibodies acting only by direct neutralization of virus and define the protective level of serum neutralizing activity in the absence of confounding variables of Fc-mediated effects. A threshold for virological protection in BAL fluid and NP secretions was estimated to be equal to or higher than 6,000 for serum neutralizing antibody titer (NT_50_), since antibody levels above these thresholds conferred full protection in 83% to 93% of challenged NHPs. This quantitative determination of a neutralizing titer as a direct mechanistic correlate of protection has implications for estimating the durability of protection conferred by passive immunization with antibodies (Loo et al., 2021) or active immunization with vaccines. The failure to achieve serum neutralizing titers above this threshold likely explains the lack of limited efficacy observed in most clinical trials of COVID-19 convalescent plasma (Begin et al., 2021; Bradfute et al., 2020; Janiaud et al., 2021). Also, this quantitative threshold for correlate of protection sheds light on the somewhat limited magnitude and durability of the humoral immunity component of protection following natural infection or immunization. Third, the studies also support a public health strategy of prophylaxis of high-risk individuals who cannot be adequately vaccinated by using administration of neutralizing mAbs instead. Engineering of the Fc region with YTE mutations to accomplish long-half extends the prophylactic efficacy of the antibodies, predicted to last for at least several months. Fourth, we also assessed IM and IV administration and found that IM administration was effective, which could allow a much easier and more practical approach to administration of these antibodies at large scale in populations at risk.

Numerous groups have reported the isolation of potently neutralizing antibodies from survivors that target the RBD of SARS-CoV-2 S protein (Brouwer et al., 2020; Cao et al., 2020; Robbiani et al., 2020; Rogers et al., 2020; Shi et al., 2020; Wec et al., 2020; Wu et al., 2020b). The studies here support the further development of a two-mAb prophylactic anti-SARS-CoV-2 cocktail (ADM03820) incorporating mAbs that target non-overlapping regions of the RBD (Zost et al., 2020a; Zost et al., 2020b). The combination of engineered antibodies possesses desirable features consistent with the objectives above, including long half-life, an effective IM formulation, accumulation at respiratory mucosa following systemic administration, and a clear mechanism of action purely through direct virus neutralization. The combination was shown effective in a stringent rhesus macaque model for SARS-CoV-2 we previously developed with high viral loads in the upper and lower respiratory tract, cellular and humoral immune responses, and pathogenic evidence of viral pneumonia (Chandrashekar et al., 2020; Yu et al., 2020). In the present study, we demonstrated that prophylactic administration of the two-mAb cocktail ADM03820 for protection against SARS-CoV-2 infection in this animal model, reducing viral loads in the upper and lower airways and accelerating virus clearance.

These antibodies include the YTE mutations in Fc region, which increase the serum half-life of the mAbs (Dall’Acqua et al., 2006; Dall’Acqua et al., 2002; Yu et al., 2017) and the LALA Fc mutations that were designed to decrease the Fc effector function by reducing interaction with Fcγ receptors (Lund et al., 1991; Wines et al., 2000; Woodle et al., 1998; Yu et al., 2017) . Studies in murine SARS-CoV-2 challenge models have demonstrated equivalently high prophylactic efficacy by potently neutralizing RBD-specific mAb variants with intact or abrogated Fc region-mediated effector functions (Winkler et al., 2021). Previous studies in a similar NHP model have shown that COV2-2381 IgG with a conventional Fc region cleared the virus infection, and no virus was observed when given at 50 mg/kg (Zost et al., 2020a). Here, the addition of YTE and LALA mutations did not appear to reduce the ability of these mAbs to clear SARS-CoV-2 infection in either the BAL fluid or NP swabs in rhesus macaques when administered three days prior to challenge.

A lower serum antibody neutralizing titer (>100) was associated with protection by vaccines in NHP SARS-CoV-2 challenge models (Corbett et al., 2020; McMahan et al., 2021; Yu et al., 2020) and in human clinical trials (Anderson et al., 2020; Jackson et al., 2020; Khoury et al., 2021) relative to the protective titer associated with mAbs (∼6,000) that we defined here. However, a similar protective titer against SARS-CoV-2 was identified in NHPs for a combination of another combination of two neutralizing human mAbs in clinical development - AZD7442 (Loo et al., 2021). Future studies are needed to determine if the lower serum neutralizing antibody protective titer for COVID-19 vaccines relative to that achieved by passive mAb transfer is due to targeting of multiple epitopes on the SARS-CoV-2 S, different anatomical distribution of antibody responses, a contribution of Fc-mediated effector functions in the polyclonal response, or complementary mechanisms of protection that are mediated by vaccine-induced T cells.

The RBD sequence is highly variable in SARS-CoV-2, which may represent a selective adaptation (Demogines et al., 2012; Frank et al., 2020; MacLean et al., 2020; Starr et al., 2020). Our approach, to use a combination of two antibodies that do not compete for the same epitope, could prevent the selection of escape mutant viruses that are likely inherent in monotherapy approaches. Recent work in the context of SARS-CoV-2 has demonstrated that combinations of two antibodies that do not compete for binding to the same region of the spike protein offer higher resistance to escape mutations while protecting animals from SARS-CoV-2 challenge (Baum et al., 2020a; Baum et al., 2020b; Weinreich et al., 2020; Zost et al., 2020a; Chen et al., 2021b).

In prior NHP studies, mAbs typically were infused via IV administration. The studies presented here demonstrate the efficacy of these antibodies either administered as a combination or alone when administered by the IM route This approach could provide a more broadly deployable route of administration for these antibodies to patients in clinical settings. In addition, the doses that were efficacious in these studies translate to very low doses in humans compared to conventional antibody therapies. The data generated in these studies provides strong evidence for the continued development of these antibodies for clinical use.

## Acknowledgements

This research was supported by a contract from the Joint Program Executive Office for Chemical, Biological, Radiological and Nuclear Defense (JPEO-CBRND) contract number W911QY-20-9-003; 20-05, the Joint Sciences and Technology Office and the Joint Program Executive Office, contract number MCDC-16-01-002 JSTO, JPEO and by DARPA grant HR0011-18-2-0001, and NIH grant R01 AI157155. J.E.C. is a recipient of the 2019 Future Insight Prize from Merck KGaA. The content is solely the responsibility of the authors and does not represent the official views of the U.S. government or other sponsors.

## MATERIALS AND METHODS

### Monoclonal antibodies

The antibody COV2-2381 and COV2-2130 sequences have been previously described (Zost et al., 2020a; Zost et al., 2020b). The antibodies were produced and purified as previously described (Tomic et al., 2019). Briefly, stably transfected CHO cells expressing either COV2-2130-YTE-LALA or COV2-2381-YTE-LALA were generated using Leap-In transposon vectors (ATUM) containing the respective antibody heavy and light chain genes and a glutamine synthetase gene as a selectable marker. Leap-In vectors were transfected into a CHO-K1 GS knockout cell line (HD-BIOP3; from Horizon Discovery) and stably transfected pools were selected using medium lacking L-glutamine. Manufacturing was performed under Good Manufacturing Practices using stably transfected pools in large scale bioreactors and antibody material was purified from harvested supernatants. The downstream processes consisted of 3 chromatography steps: 3) viral inactivation, 2) filtered viral reduction (Planova), and 3) an ultrafiltration step to concentrate the product to the appropriate g/L. Both individual antibodies and the combination were generated as cGMP-grade drug substance and drug product materials, were provided at a concentration of 52 mg/mL and were stored at -80°C until day of administration. On the day of administration, the stock vials were thawed at room temperature (RT) and gently inverted 6 to 10 times to mix the contents. After thawing, the vials were stored at RT until use. Based on individual animal weights and dose required, the purified antibody stock for each NHP was diluted to 1 mL in 0.9% normal saline diluent (Baxter) for IM injections and 10 mL in the same diluent for IV infusions. IM injections were delivered bilaterally in the upper quadriceps at 0.5 mL/quadriceps. IV infusions were performed at a rate of 1 to 2 mL/min over 5 to 10 min/animal for a total of 10 mL infused per animal.

### Animal studies

All animals were maintained at Bioqual, Inc. (Rockville, MD) which is fully accredited by the Association for Assessment and Accreditation of Laboratory Animal Care International (AAALAC) and approved by the Office of Laboratory Animal Welfare (NIH/PHS assurance number D16-00052). Studies were conducted in compliance with all relevant local, state, and federal regulations and were approved by the Bioqual Institutional Animal Care and Use Committee (IACUC). Cynomolgus monkeys (*Macaca fascicularis*) (2.2 – 5.8 kg body weight; 6 to 12 years old) were mixed male and female and randomly assigned to groups. In Study 1 (n=3/group), experimental animals received the ADM03820 cocktail of COV2-2130-YTE-LALA and COV2-2381-YTE-LALA at either 11.7 mg/kg IM or 31.3 mg/kg IV and were followed for 12 weeks for antibody pharmacokinetics only without any SARS-CoV-2 challenge. In Study 2 (n=4/group), sham control animals received no mAb while 12 experimental animals were administered the ADM03820 cocktail at varying doses and administration routes three days before challenge as described in **Figure 2**. Animals then were challenged with 10^5^ TCID_50_ SARS-CoV-2 USA-WA1/2020. These doses were administered as 0.5 mL per nare intranasally and 1 mL intratracheally on day 0. In Study 3, four sham-treated controls received no mAb while 12 experimental animals (n=3/group) were administered three days prior to challenge with either COV2-2130-YTE-LALA or COV2-2381-YTE-LALA separately at varying doses and administration routes as described in **Figure 3A**. Animals were then challenged with 10^5^ TCID_50_ SARS-CoV-2 similarly as in the first study. In Study 4 (n=3/group), sham control animals received no mAb while experimental animals were administered the ADM03820 cocktail IM at varying low doses three days before challenges performed similarly to studies 2 and 3 (**Figure 2,3**). Macaques in all four studies were monitored daily with an internal scoring protocol approved by the IACUC. These studies were not blinded.

### Viruses

The SARS-CoV-2 USA-WA1/2020 strain was obtained from BEI Resource (NR-52281; Lot #7003175). The viral stocks were expanded using Vero E6 cells and harvested on day 5 following inoculation. To confirm the viral identity, complete genome sequencing was performed and was shown to be 100% identical to the parent virus sequence. The D614G virus was produced by introducing the mutation into an infectious clone of WA1/2020, and the B.1.351 and B.1.1.28 spike genes were cloned into the WA1/2020 infectious clone to produce Wash-B.1.351 and Wash-B.1.1.28 chimeric viruses, as described previously (Chen et al., 2021a). B.1.1.7 and B.1.617.2 were isolated from infected individuals. D614G, Wash-B.1.351, Wash-B.1.1.28, B.1.1.7, and B.1.617.2 viruses were propagated on Vero-TMPRSS2 cells and subjected to deep sequencing.

### Quantification of circulating human mAbs and serum neutralization activity

The quantification of infused/injected human SARS-CoV-2 mAbs in NHP serum at multiple time points was performed as previously described [20]. Additionally, the serum neutralization activities of infused or injected mAbs were also monitored at the same time points using a pseudovirus neutralization assay as previously described [24, 25].

### BAL and NP swab collection

Collection of mucosal secretions was performed on sedated NHPs using cotton swabs (COPAN flocked swab) or nasosorption FX-I devices (Hunt Developments Ltd.). The swabs were inserted into the nasal cavity and rotated gently. Following collection, the swabs were placed into a collection vial containing 1 mL of phosphate buffered saline (PBS). All vials were stored at ≤-70°C until viral load testing (or antibody quantification if required).

The bronchoalveolar lavage (BAL) collection procedure was performed on anesthetized animals by the “chair method”. In brief, each animal was placed in dorsal recumbency in a chair channel and a red rubber feeding tube inserted into the trachea via a laryngoscope during inspiration. A total of 10 mL PBS was flushed through the tube and the volume instilled and recovered from each animal recorded. The collected BAL samples were placed immediately onto wet ice and processed for isolation of fluid by centrifugation at 4°C followed by supernatant removal. BAL aliquots were stored at ≤-70°C until viral load testing (or antibody quantification if required).

### Focus reduction neutralization test

Serial dilutions of mAbs were incubated with 10^2^ FFU of different strains or variants of SARS-CoV-2 for 1 h at 37 °C. Antibody–virus complexes were added to Vero-TMPRSS2 cell monolayers in 96-well plates and incubated at 37 °C for 1 h. Subsequently, cells were overlaid with 1% (w/v) methylcellulose in MEM. Plates were collected 30 h later by removing overlays and fixed with 4% PFA in PBS for 20 min at room temperature. Plates were washed and sequentially incubated with an oligoclonal pool of SARS2-2, SARS2-11, SARS2-16, SARS2-31, SARS2-38, SARS2-57 and SARS2-71 anti-S (VanBlargan et al., 2021) antibodies and HRP-conjugated goat anti-mouse IgG (Sigma, 12-349) in PBS supplemented with 0.1% saponin and 0.1% bovine serum albumin. SARS-CoV-2-infected cell foci were visualized using TrueBlue peroxidase substrate (KPL) and quantitated on an ImmunoSpot microanalyzer (Cellular Technologies).

### Subgenomic mRNA assay

The subgenomic mRNA of SARS-CoV-2 was assessed by RT-PCR as previously described (Chandrashekar et al., 2020; Wolfel et al., 2020; Yu et al., 2020). The standard curve is based on the SARS-CoV-2 E gene. Prior to PCR, cDNA was generated from each animal using Superscript III VILO (Invitrogen) according to the manufacturer’s instructions. Using the sequences targeting the E gene mRNA, a TaqMan custom gene expression assay (Thermo Fisher Scientific) was designed (Wolfel et al., 2020) and reactions were carried out using a QuantStudio 6 and 7 Flex Real-Time PCR system (Applied Biosystems) according to the manufacturer’s instructions. Standard curves were generated to calculate sgRNA/mL or per swab. Viral load for each timepoint tested per NHP was reported as the average of two replicates. The sensitivity of this assay was 50 copies per mL of BAL or per swab.

### Quantification and statistical analysis

The average change in viral load (log_10_ sgRNA copies/mL or swab) was assessed from day 1 to day 14 (Study 2), or from day 1 to day 10 (Study 3 and 4). The time-weighted average (TWA) values for the change of sgRNA viral load in BAL or NP from day 1 to day 10 after viral challenge were calculated as the area under the curve (AUC) of the change in viral load in Prism (version 9.1.2; GraphPad) and then divided by 10 as described previously (Baum et al., 2020a) (**Table S1**). The TWA values of each treatment group were compared to those of the sham group using Welch’s t-test. The significance level alpha of 10% was pre-specified, and estimated P-values are indicated in the figures. TWA threshold was set up to ≤ 0.3 for full protection, ≤ 0.51 (the lower sham point) for partial protection, and > 0.51 for no protection in BAL, and ≤ 0.3 for full protection, < 1.638 (the lower sham point), for partial protection, and > 1.638 for no protection in NP. To estimate protective antibody concentration or neutralizing titer in serum, the optimal thresholds that maximizes the sum of sensitivity and specificity for full protection were calculated and reported in **Table S2**. Sensitivity is the proportion above the threshold in the fully-protected subjects, and specificity is the proportion below the threshold in partially- or non-protected subjects. The fitting curves and confidence intervals to visualize the relationship between TWA and antibody levels were estimated using Lowess curve smoothing method using ggplot2 in R software. The other data visualization was performed using Prism software.

