## Supplementary material for "A combination of two human neutralizing antibodies prevents SARS-CoV-2 infection in rhesus macaques": Table S1 and Table S2

**Table S1. Measurements to determine thresholds for antibody-mediated protection against viral challenge in BAL or NP sites in NHP. Related to Figure 8.**

|  | | | | | **BAL** | | | **NP swab** | | |
| --- | --- | --- | --- | --- | --- | --- | --- | --- | --- | --- |
| **Study** | **Treatment** | **NHP ID** | **Day 0 Ab concentration (log10μg/mL)*** | **Day 0 Ab log10NT50**** | **Day 0-10 area under the curve (AUC) sgRNA level** | **Day 0-10 area under the curve (AUC) sgRNA level minus AUC of LOD***** | **Time-weighted average (TWA) viral sgRNA load** | **Day 0-10 area under the curve (AUC) sgRNA level** | **Day 0-10 area under the curve (AUC) sgRNA level minus AUC of LOD***** | **Time-weighted average (TWA) viral sgRNA load** |
| **2** | **Sham** | V293 | -2.00 | 1.30 | 22.1 | 5.10 | 0.51 | 39.72 | 22.70 | 2.27 |
|  |  | V294 | -2.00 | 1.30 | 34.75 | 17.80 | 1.78 | 41.83 | 24.80 | 2.48 |
|  |  | V280 | -2.00 | 1.30 | 25.75 | 8.80 | 0.88 | 35.58 | 18.60 | 1.86 |
|  |  | V284 | -2.00 | 1.30 | 27.01 | 10.00 | 1.00 | 38.72 | 21.70 | 2.17 |
|  | **3.9 mg/kg I.M.** | V313 | 1.18 | 4.33 | 16.99 | 0.00 | 0.00 | 30.29 | 13.30 | 1.33 |
|  |  | V318 | 1.48 | 4.31 | 16.99 | 0.00 | 0.00 | 18.99 | 2.00 | 0.20 |
|  |  | V286 | 1.13 | 3.92 | 18.2 | 1.20 | 0.12 | 19.09 | 2.10 | 0.21 |
|  |  | V290 | 1.34 | 3.92 | 16.99 | 0.00 | 0.00 | 19.39 | 2.40 | 0.24 |
|  | **11.7 mg/kg I.M.** | V324 | 1.70 | 4.21 | 16.99 | 0.00 | 0.00 | 18.81 | 1.80 | 0.18 |
|  |  | V291 | 1.79 | 5.02 | 17.83 | 0.80 | 0.08 | 16.99 | 0.00 | 0.00 |
|  |  | V299 | 1.92 | 5.35 | 18.15 | 1.20 | 0.12 | 19.73 | 2.70 | 0.27 |
|  |  | V303 | 1.96 | 4.59 | 16.99 | 0.00 | 0.00 | 19.11 | 2.10 | 0.21 |
|  | **31.3 mg/kg I.V.** | V331 | 2.63 | 5.69 | 16.99 | 0.00 | 0.00 | 19.05 | 2.10 | 0.21 |
|  |  | V308 | 2.36 | 5.64 | 16.99 | 0.00 | 0.00 | 19.27 | 2.30 | 0.23 |
|  |  | V314 | 2.59 | 5.74 | 16.99 | 0.00 | 0.00 | 18.51 | 1.50 | 0.15 |
|  |  | V315 | 2.55 | 5.27 | 16.99 | 0.00 | 0.00 | 16.99 | 0.00 | 0.00 |
| **3** | **Sham** | V298 | -2.00 | 1.30 | 34.76 | 17.80 | 1.78 | 41.77 | 24.80 | 2.48 |
|  |  | V302 | -2.00 | 1.30 | 27.36 | 10.40 | 1.04 | 35.32 | 18.30 | 1.83 |
|  |  | V319 | -2.00 | 1.30 | 38.39 | 21.40 | 2.14 | 41.07 | 24.10 | 2.41 |
|  |  | V326 | -2.00 | 1.30 | 31.51 | 14.50 | 1.45 | 44.81 | 27.80 | 2.78 |
|  | **1.95 mg/kg I.M.** | V306 | 1.17 | 3.50 | 18.37 | 1.40 | 0.14 | 24.95 | 8.0 | 0.80 |
|  |  | V307 | 1.33 | 4.0 | 16.99 | 0.00 | 0.00 | 35.66 | 18.70 | 1.87 |
|  |  | V296 | 1.45 | 4.30 | 18.83 | 1.80 | 0.18 | 16.99 | 0.00 | 0.00 |
|  | **15.65 mg/kg I.V.** | V309 | 2.33 | 4.60 | 18.53 | 1.50 | 0.15 | 19.2 | 2.20 | 0.22 |
|  |  | V311 | 2.24 | 4.40 | 17.99 | 1.00 | 0.10 | 18.53 | 1.50 | 0.15 |
|  |  | V300 | 2.34 | 4.70 | 18.93 | 1.90 | 0.19 | 19.42 | 2.40 | 0.24 |
|  | **1.95 mg/kg I.M.** | V316 | 1.11 | 2.50 | 16.99 | 0.00 | 0.00 | 19.36 | 2.40 | 0.24 |
|  |  | V301 | 1.13 | 2.60 | 24.89 | 7.90 | 0.79 | 40.82 | 23.80 | 2.38 |
|  |  | V304 | 1.17 | 2.30 | 18.96 | 2.00 | 0.20 | 22.85 | 5.90 | 0.59 |
|  | **15.65 mg/kg I.V.** | V317 | 2.19 | 3.80 | 16.99 | 0.00 | 0.00 | 19.26 | 2.30 | 0.23 |
|  |  | V310 | 2.31 | 3.80 | 18.68 | 1.70 | 0.17 | 18.93 | 1.90 | 0.19 |
|  |  | V312 | 2.28 | 3.40 | 18.65 | 1.70 | 0.17 | 18.68 | 1.70 | 0.17 |
| **4** | **Sham** | GA945H | -2.00 | 1.30 | 27.37 | 10.40 | 1.04 | 33.37 | 16.40 | 1.64 |
|  |  | V320 | -2.00 | 1.30 | 28.36 | 11.4 | 1.14 | 36.92 | 19.90 | 1.99 |
|  |  | V305 | -2.00 | 1.30 | 32.52 | 15.50 | 1.55 | 37.89 | 20.90 | 2.09 |
|  | **3.91 mg/kg I.M.** | V328 | 1.59 | 3.86 | 21.56 | 4.60 | 0.46 | 26.04 | 9.10 | 0.91 |
|  |  | V289 | 1.53 | 3.82 | 18.85 | 1.90 | 0.19 | 19.19 | 2.20 | 0.22 |
|  |  | V295 | 1.61 | 4.30 | 21.2 | 4.20 | 0.42 | 19.4 | 2.40 | 0.24 |
|  | **1.95 mg/kg I.M.** | V281 | 1.13 | 3.75 | 20.27 | 3.30 | 0.33 | 23.92 | 6.90 | 0.69 |
|  |  | V321 | 1.11 | 4.03 | 22.67 | 5.70 | 0.57 | 29.41 | 12.40 | 1.24 |
|  |  | V288 | 1.14 | 3.96 | 20.72 | 3.70 | 0.37 | 18.94 | 2.00 | 0.20 |
|  | **0.98 mg/kg I.M.** | V329 | 0.85 | 3.31 | 16.99 | 0.00 | 0.00 | 35.89 | 18.90 | 1.89 |
|  |  | V330 | 0.90 | 2.90 | 18.24 | 1.30 | 0.13 | 21.13 | 4.10 | 0.41 |
|  |  | V285 | 0.83 | 3.01 | 21.02 | 4.00 | 0.40 | 29.58 | 12.60 | 1.26 |
|  | **0.49 mg/kg I.M.** | V323 | 0.43 | 2.58 | 23.46 | 6.50 | 0.65 | 34.45 | 17.50 | 1.75 |
|  |  | V327 | 0.60 | 3.40 | 20.59 | 3.60 | 0.36 | 35.04 | 18.10 | 1.81 |
|  |  | V282 | 0.62 | 3.01 | 21.12 | 4.10 | 0.41 | 27.65 | 10.70 | 1.07 |

* Indicates LOD of antibody concentration measurement in serum with the value that is equal to 10 ng/mL (designated to zero in **Figure 8A**); ** indicates LOD of antibody neutralizing titer measurement in serum with the value that is equal to 1.3log10 NT_50_ (designated to zero in **Figure 8A**); ***AUC LOD value was estimated for the curve from day 0 to 10 with viral sgRNA load below the detection limit on each timepoint and was equal to 16.99.

**Table S2.** **Estimated specificity, sensitivity, and cut-off values for protective mAb concentration or titer in NHP serum. Related to Figure 8.**

| **Site** | **BAL** | | | **NP swap** | | |
| --- | --- | --- | --- | --- | --- | --- |
| **Measurement** | **Sensitivity** | **Specificity** | **Cut-off*** | **Sensitivity** | **Specificity** | **Cut-off*** |
| **Antibody concentration** | 0.85 | 0.90 | 1.17 | 0.86 | 0.96 | 1.34 |
| **Antibody neutralizing titer** | 0.76 | 0.81 | 3.8 | 0.91 | 0.84 | 3.8 |

* log_10_μg/mL for mAb concentration and log_10_ NT_50_ for neutralizing titer measurements.
